# *Fusarium oxysporum* Effector Clustering version 2 (FoEC2): an updated pipeline to infer host range

**DOI:** 10.1101/2022.08.08.503139

**Authors:** Megan A. Brenes Guallar, Like Fokkens, Martijn Rep, Lidija Berke, Peter van Dam

## Abstract

The fungus *Fusarium oxysporum* is infamous for its devastating effects on economically important crops worldwide. *F. oxysporum* isolates are grouped into *formae speciales* based on their ability to cause disease on different hosts. Assigning *F. oxysporum* strains to *formae speciales* using non-experimental procedures has proven to be challenging due to their genetic heterogeneity and polyphyletic nature. However, genetically diverse isolates of the same *forma specialis* encode similar repertoires of effectors, proteins that are secreted by the fungus and contribute to the establishment of compatibility with the host. Based on this observation, we previously designed the *F. oxysporum* Effector Clustering (FoEC) pipeline which is able to classify *F. oxysporum* strains by *forma specialis* based on hierarchical clustering of the presence of predicted putative effector sequences, solely using genome assemblies as input. Here we present the updated FoEC2 pipeline which is more user friendly, customizable and, due to multithreading, has improved scalability. It is designed as a Snakemake pipeline and incorporates a new interactive visualization app. We showcase FoEC2 by clustering 537 publicly available *F. oxysporum* genomes and further analysis of putative effector families as multiple sequence alignments. We confirm classification of isolates into *formae speciales* and are able to further identify their subtypes. The pipeline is available on github: https://github.com/pvdam3/FoEC2.

## 2 Introduction

The cosmopolitan *Fusarium oxysporum* species complex (FOSC) includes well known plant pathogens that cause diseases in a broad range of hosts such as tomato, onion and banana (Edel-Hermann et al., 2019). Pathogenic *F. oxysporum* strains are a significant threat to crop production and cause devastating decreases in yield and economic losses (Cook et al., 2015; Panno et al., 2021).

*F. oxysporum* strains are grouped into a *forma specialis* (f. sp.) depending on the hosts they are capable of infecting. More than a hundred *formae speciales* (ff. spp.) have been documented to date, varying in their host range from a single species to multiple genera (Edel-Hermann et al., 2019). However, strains grouped into the same *forma specialis* are not necessarily phylogenetically closely related (Baayen et al., 2000). The polyphyletic nature of host range in FOSC has hampered the determination of *forma specialis* of uncharacterized *F. oxysporum* strains using conserved gene sequences. Experimental methods can be used and remain the gold standard for host determination, but are less favorable due to their time- and labor-intensive nature.

Effector genes encode small secreted proteins that enable host colonization, for instance by suppressing host immunity. The genome of *F. oxysporum* f. sp. *lycopersici* encodes 14 such effectors named Secreted In Xylem (*SIX1* - *SIX14*) which have been identified in xylem sap of infected tomato plants (Rep et al., 2004; Houterman et al., 2004; Ma et al., 2010; Schmidt et al., 2013). Genomic analyses uncovered that the promoter regions of *SIX* genes frequently contain miniature impala (mimp) transposable elements (TEs) (Schmidt et al., 2013). While mimps do not seem to be directly involved in the transcriptional regulation of *SIX* genes (Schmidt et al., 2013), their presence is correlated to putative horizontal gene transfer events involving effector genes (van Dam et al., 2017b). Horizontal chromosome transfer (HCT) of accessory chromosomes or chromosome fragments is believed to be one of the driving factors facilitating the spread of host specificity between isolates (Ma et al., 2010, Li et al., 2020).

The *F. oxysporum* Effector Clustering (FoEC) pipeline (van Dam et al., 2016) is a computational method to predict candidate effector genes and cluster these based on their presence/absence patterns. Traditional gene prediction methods have trouble detecting effector genes due to their short sequence, fast evolutionary rate and localization in complex genomic regions (Gibriel et al., 2016). To circumvent these, the FoEC pipeline exploits the presence of (partial) mimps in the promoter regions of *F. oxysporum* effector genes, reducing the search space significantly. Mimps are easily identified by their terminal inverted repeats (TIRs), which can be found by their consensus sequence (Bergemann et al., 2008). When using this pipeline, *F. oxysporum* strains that belong to the same *forma specialis* are typically grouped together, solidifying the hypothesis that the effector repertoire of a *F. oxysporum* genome plays a role in determining the host range of a strain. FoEC has been used to classify uncharacterized *F. oxysporum* strains into potential *formae speciales* based on presence/absence patterns of putative effector genes (Urbaniak et al., 2019; Constantin et al., 2021; Sabahi et al., 2021), considerably narrowing down the number of hosts to test in experimental procedures.

Here we present the updated FoEC2 pipeline that improves upon both usability and functionality. FoEC2 is implemented in Python3 and based in Snakemake (Mölder et al., 2021) to help manage external tools and internal scripts, as well as improve scalability with multithreading. It uses hidden Markov models (HMM) profile search instead of BLAST to increase the sensitivity of searches, and R Shiny to visualize presence/absence patterns of putative effectors and their multiple sequence alignments (MSAs). We benchmark and demonstrate FoEC2 on a dataset of 537 currently available

FOSC genomes and demonstrate that previously uncharacterized isolates can be assigned to their most probable *formae speciales*.

## 3 Materials and Methods

### 3.1 Implementation

FoEC2 is available on github: https://github.com/pvdam3/FoEC2. It is based on Python3 and Snakemake (v. 6.15.1) and uses conda environments for installation of dependencies. A minimal command to run FoEC2 requires a directory containing *F. oxysporum* genomes as input. Optional input files include genome annotations to supplement the predictions made by FoEC2 and known effector sequences to skip the prediction part of the pipeline.

Several configuration files are used in the Snakemake pipeline. The genome configuration file contains the paths to all the input genomes along with their labels (by default, the file name), as well as the paths to their annotation files if provided. A bash script is used to facilitate writing this file, taking the provided input files and producing the genome configuration file. Filtering thresholds such as ORF length, number of cysteines, three- and six-frame translation and size of search region can also be modified in the main configuration file.

#### 3.1.1 Identification of putative effectors

Mimp TIRs are identified by a regular expression with their consensus sequence (“TT[TA]TTGCNNCCCACTGNN”) (Bergemann et al., 2008, van Dam et al., 2017b). Open reading frames (ORFs) are located in either three- or six-frame translation mode using a custom Python3 script. In three-frame mode, which is used throughout this manuscript, ORFs must be downstream of a TIR and point away from the TIR. Six-frame mode is included to locate putative effectors anywhere surrounding the TE, a feature which may be valuable when searching for other types of genomic elements associated with effectors.

SignalP (Almagro Armenteros et al., 2019) is used to detect signal peptides in translated ORFs (‘-format short -gff3 -batch 10000’). FoEC2 supports SignalP versions 4.1 and 5; version 5.0b was used for the analysis described below. SignalP is under an academic software license and is therefore the only tool which requires manual installation. AUGUSTUS (Stanke et al., 2006; v3.4.0; parameters ‘--species=fusarium --genemodel=complete --noInFrameStop=true --predictionStart=X -- predictionEnd=X --strand=X’) is used to update the ORFs with signal peptides. In case of overlap, the gene model by AUGUSTUS is retained. The resulting ORFs are filtered on length and presence of cysteines (defaults: 20aa ≤ size ≤ 600aa; 0 ≤ cysteines) and are considered putative effectors.

#### 3.1.2 Identification of effector clusters

Clustering on all putative effector protein sequences is performed using Diamond v2.0.13 (Buchfink et al., 2015) with Diamond BLASTP. Next, MCL v14.137 (Enright et al., 2002; Van Dongen et al., 2012; i = 1.2) creates clusters of putative effectors.

#### 3.1.3 Presence/absence variation

An MSA of each gene cluster was generated with MAFFT v7.490 (Katoh et al., 2013; default parameters), followed by constructing a hidden Markov model (HMM) profile with HMMER v3.3.2 (Eddy, 2011). The putative effector HMM profiles queries for the input genomes using HMMER’s nhmmer function (Wheeler, Eddy, 2013). A hit is valid if the E-value (default = 10E-10) and query coverage (default = 80%) meet the provided thresholds.

#### 3.1.4 Clustering and visualization

R Shiny was used to visualize results with R v4.0.4 (R Core Team, 2020). The distance and clustering methods for both genomes (rows) and putative effectors (columns) can be adjusted by the user and are set at ‘binary’ distance and ‘average’ clustering by default. The following packages were used: shiny v1.7.1, shinythemes v1.2.0, dendextend v1.15.2, RColorBrewer v1.1-2, pals v1.7, pheatmap v1.0.12, phylogram v2.1.0, DT v0.20, rhandsontable v0.3.8, msaR v0.6.0 (for the BioJS MSA viewer (Yachdav et al., 2016).

### 3.2 Datasets

All publicly available FOSC genomes (as of January 18, 2022) were obtained from NCBI (Supplementary Table S1). Duplicates of the *F. oxysporum* strain 4287 (GCA_003315725.1, GCA_001703185.1, GCA_001703175.2, GCF_000149955.1) and f. sp. *melonis* NRRL_26406 (GCA_002318975.1) were removed, as were the samples VCG0125, VCG0120 and VCG01220 (GCA_016802195.1, GCA_016802205.1, GCA_016802225.1) as they were *in silico* MISAG hybrids where sequencing data from f. sp. *cubense* was imputed using Fol4287 as a reference. Due to low quality of some genomes, the completeness of each accession was determined using BUSCO (Manni et al., 2021; v. 5.30; using lineage ‘hypocreales_odb10’, 4494 genes). Only genomes with a BUSCO completeness score of 70% or higher were used for further analysis, leaving a total of 537 *F. oxysporum* samples. The *F. verticillioides* genome (GCF_000149555) was added as an outgroup. An overview of all removed samples can be seen in Supplementary Table S2. To benchmark the FoEC2 pipeline, the dataset with 59 isolates described by van Dam et al. (2016) (Supplementary Table S3) was used, as well as the FoEC pipeline available at https://github.com/pvdam3/FoEC (default settings).

The sequences of *SIX* genes were obtained from van Dam et al. (2016). *SIX* genes (*SIX1*-*SIX14*) were identified among the final putative effectors with blastn (v. 2.12.0+; task = blastn-short; E-value threshold 1E-5).

### 3.3 Phylogeny construction

A phylogeny was constructed using the set of 537 publicly available *F. oxysporum* genomes (Supplementary Table S1), as well the *F. verticillioides* 7600 genome (GCF_000149555). This phylogeny was based on single copy BUSCO genes found in each genome using the ‘hypocreales_odb10’ lineage. BUSCO genes found in at least 98% of all genomes were used as input for MAFFT v7.490 to generate MSAs based on translated amino acid sequences. The resulting MSAs were trimmed with trimAl v1.4 (Capella-Gutiérrez et al., 2009; parameter ‘gappyout’) and concatenated into a single FASTA file, with a total length of 1,444,221 aligned amino acid positions. The concatenated FASTA file was then used as input for IQ-TREE v2.2.0 (Nguyen et al., 2015; parameters ‘-m JTT+F+R2 -bb 1000’) to create a phylogeny.

## 4 Results

### 4.1 The FoEC2 pipeline

The FoEC2 pipeline, like its predecessor, is divided into four main parts (Figure 1): (i) identification of putative effectors, (ii) identification of effector families, (iii) establishment of presence/absence variation, and (iv) clustering and visualization. The first three parts are executed by a Snakemake pipeline, which produces the output needed for the fourth part which performs clustering and visualization of presence/absence patterns.

**Figure 1:**
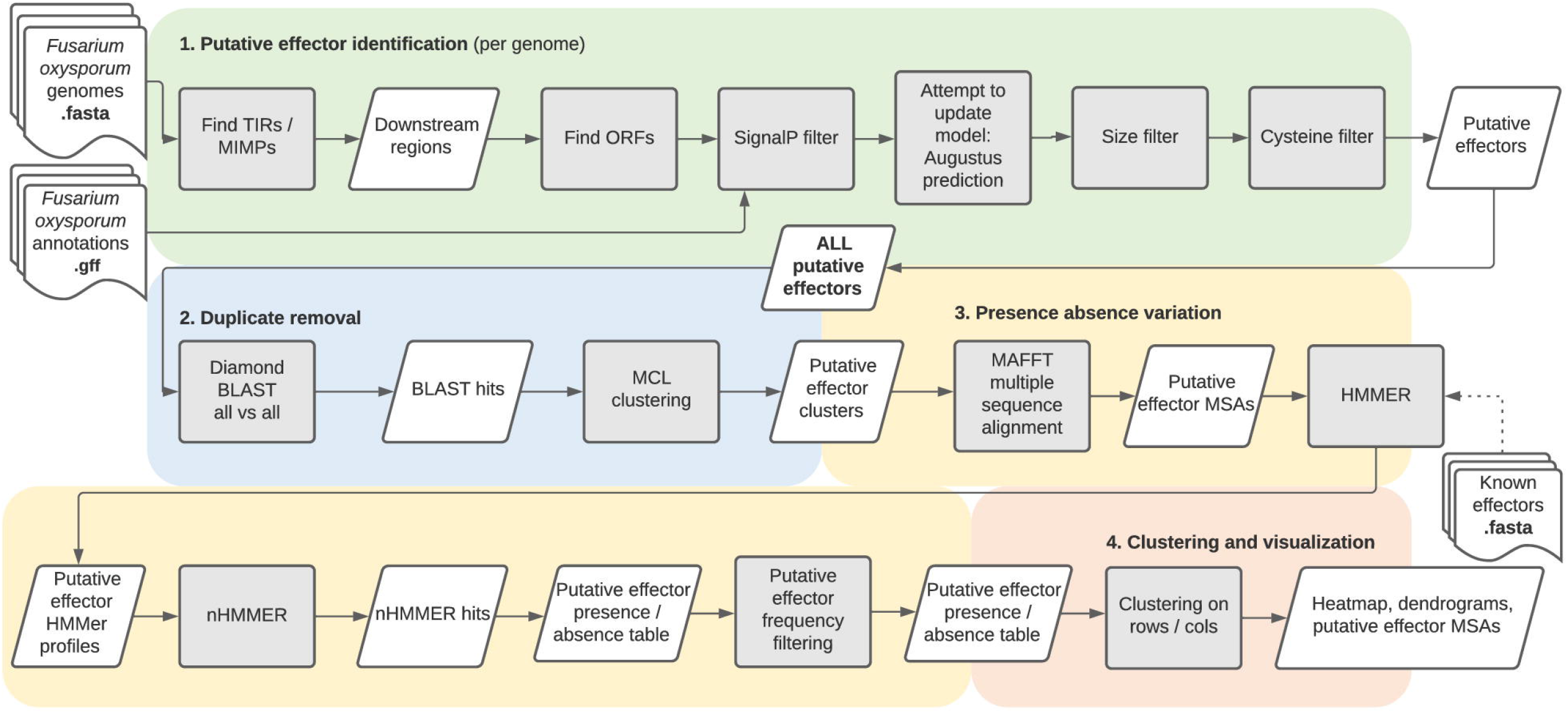
An overview of the FoEC2 pipeline.

First, mimp TIRs are searched for throughout each genome using their consensus sequence. The detection of mimp TIRs can lead to the identification of complete mimps, in which two mimp TIRs in opposite direction are found within 400 bp of one another, and incomplete mimps (solo-mimp TIRs).

The next step searches for ORFs in the vicinity (<2500 bp) of the previously identified (in)complete mimps. Translation of the input genome assembly is performed in three-frame mode, where ORFs must be downstream of a TIR and point away from that TIR. ORFs identified encode a methionine followed by sequence of at least 20 amino acids and a stop codon. Next, genes from the (optional) user-provided annotations are merged with the ORFs detected by the pipeline if they are located in the ORF search regions. In the case of overlapping genes/ORFs from both sources, the user-provided gene takes precedence over the pipeline ORF prediction. Because effector proteins are secreted, SignalP (Almagro Armenteros et al., 2019) is then used select ORFs with a signal peptide. These genomic regions, containing ORFs with a predicted signal peptide, are used as input sequences for AUGUSTUS (Stanke et al., 2006) to predict a gene model. If no gene model is predicted by AUGUSTUS, the original ORF (without introns) is taken along as is. These ORFs and gene models are then optionally filtered by size and cysteine content to decrease the number of false positives. The sequences remaining after these filtering steps are considered to be putative effectors and are available as both nucleotide and protein FASTA files.

This first part of the pipeline predicts putative effectors for each genome. Multiple isolates can share the same or very similar effectors and one isolate can have multiple copies of an effector. The pipeline removes this redundancy by clustering homologous effectors into effector families based on sequence similarity. To determine the presence of putative effector families in the initial set of genomes, each cluster is represented by an HMM profile. Searching with an HMM profile instead of a single representative sequence is one of the major differences with the original FoEC pipeline. It potentially increases sensitivity as it accounts for sequence diversity. The putative effector hits are then tallied per genome, resulting in a presence/absence variation table for each putative effector family and each genome. The final filtering step removes putative effectors which have over 20 hits in any given genome, or which have 10 or more hits on average across all genomes, as these are not likely to be effector genes but more probably (part of) transposable elements.

Results are visualized in an R Shiny app (Supplementary Figure S1), showing a customizable heatmap depicting clusters, dendrograms for genomes and putative effectors, and MSAs of putative effector clusters. Metadata for both genomes and putative effectors may also be uploaded, edited and again downloaded. Several aesthetic modifications can be made and data can be downloaded.

### 4.2 Comparison to FoEC

First, the results of FoEC2 were compared to the FoEC pipeline using the 59 isolates analyzed by van Dam et al. (2016). These include isolates of the ff. spp. *niveum, cucumerinum, radicis-cucumerinum, melonis, lycopersici, conglutinans, radicis-lycopersici, cubense, pisi* and *vasinfectum*. This analysis was completed after 2.2 hours on 50 CPUs. FoEC finished more rapidly on this small number of genomes (1.5 hours on similar infrastructure).

The FoEC2 pipeline resulted in 227 putative effector clusters (Supplementary Figure S2), compared to the 215 clusters found by FoEC (van Dam et al., 2016)). Next, the presence/absence profiles of effector families were used to cluster the isolates. On the original dataset of 59 genomes, putative effector patterns resulted in clusters that consistently grouped samples from the same *forma specialis* together. *Formae speciales* were mostly contained in single clusters, the exception being *F. oxysporum* f. sp. *cucumerinum*, which was split into two clusters, as was observed in FoEC (van Dam et al., 2016).

### 4.3 Effector clustering patterns: filling in the host specificity gaps

To test the capability of FoEC2, the pipeline was run on all 537 publicly available *F. oxysporum* genomes meeting the quality thresholds described in the Materials and Methods (Figure 2, Supplementary Table S1). The genome of *F. verticillioides* was added as an outgroup. In total, 222 out of 537 genomes contained a *forma specialis* annotation in their description. This analysis was completed after 29.7 hours on 50 CPUs.

**Figure 2:**
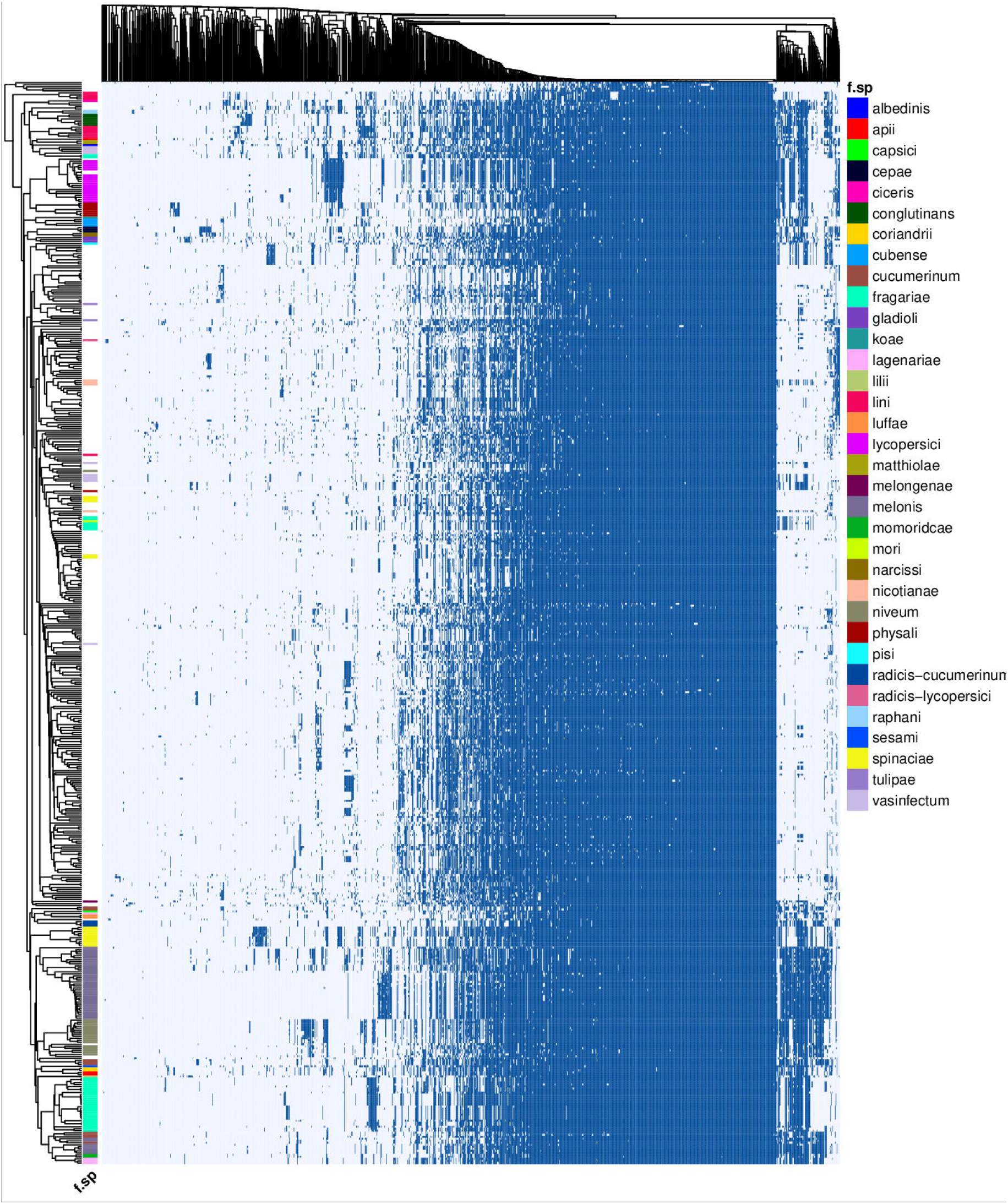
Presence (dark blue) / absence (light blue) variation of 1,283 putative effectors (columns) across 538 *Fusarium* genomes (rows). Genomes are annotated by *forma specialis* when possible. Presence is determined by nhmmer, given the thresholds E-value ≤ 10E-10, query coverage ≥ 80%. Clusters were generated with a binary distance matrix and average clustering.

In total, 1,251 putative effector families were identified. Hierarchical clustering of the presence/absence patterns of these families in the 537 *F. oxysporum* genomes (Figure 2) shows that in many cases (e.g., ff. spp. *lycopersici, niveum, conglutinans, raphani, cubense, physalis*, and others), these groups encompass all representatives of a single *forma specialis*. This approach thus allows for inference of the pathogenic potential of a *F. oxysporum* genome for which no *forma specialis* is described. For example, assemblies RBG7064 (GCA_009298205.1) and RBG7070 (GCA_009298435.1) are not assigned to a *forma specialis* (Figure 3A). However, they are found in a cluster of genomes annotated as f. sp. *niveum* and their putative effector presence pattern matches that of various *niveum* isolates. The metadata available for these two assemblies show that the samples were collected from *Citrullus lanatus* (watermelon). This means that these assemblies are likely *F. oxysporum* f. sp. *niveum*. Another example is assemblies VPRI32264 (GCA_009298805.1), VPRI16234 (GCA_009298505.1) and VPRI11681 (GCA_009298875.1) that do not have a *forma specialis* annotation, but group inside the *F. oxysporum* f. sp. *lycopersici* cluster (Figure 3B). While all three samples originated from *Solanum* samples, only VPRI32264 and VPRI11681 were collected from *Solanum lycopersicum* (tomato), whereas VPRI16234 was collected from *Solanum tuberosum* (potato). This could be an indication of an opportunistic infection of potato by a *F. oxysporum* f. sp. *lycopersici* isolate. A similar situation is shown in Figure 3C; nine isolates collected from *Pisum sativum* plants and one collected from soil (RBG5783, GCA_009297405.1) cluster closely together with the *F. oxysporum* f. sp. *pisi* HDV247 isolate originally sequenced by the Broad institute, indicative of being the same *forma specialis*. Indeed, they all possess a similar set of ‘*pisi*’ putative effectors (arrow, Figure 3C). The isolates mentioned for these three example cases were all sequenced and assembled in a study on phylogenetic relationships between Australian *F. oxysporum* isolates (Achari et al., 2020).

**Figure 3:**
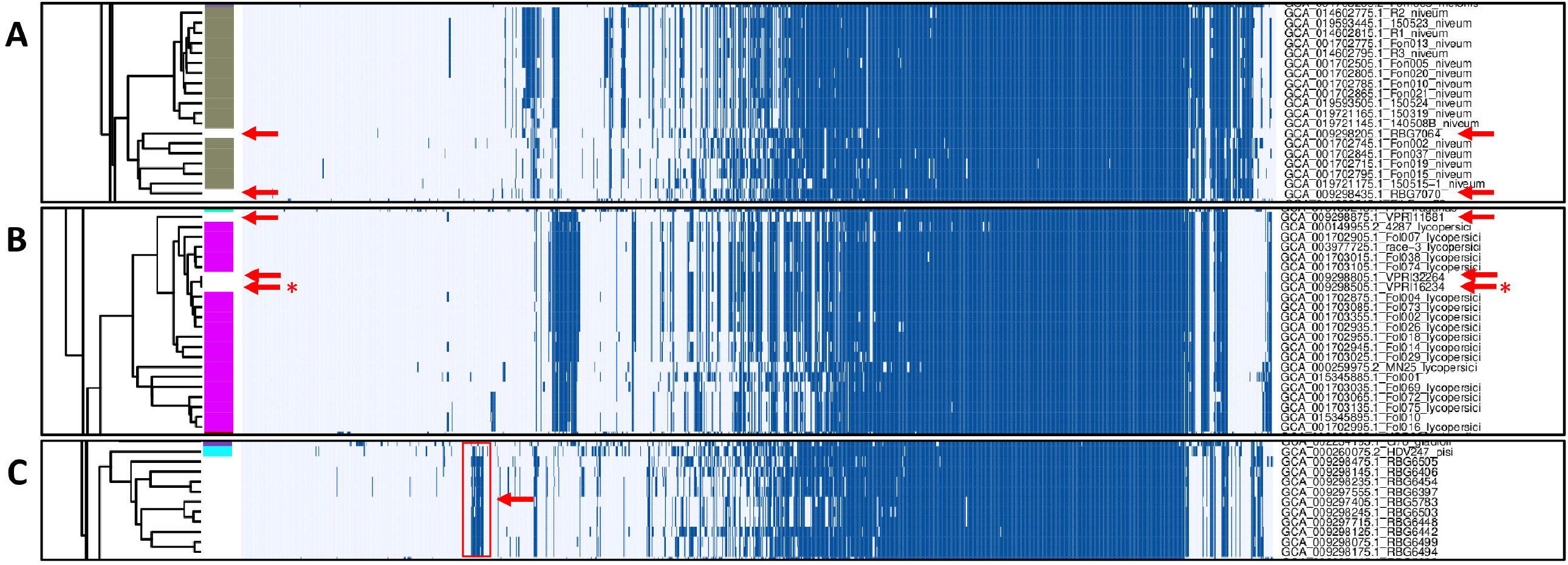
Zoomed-in examples of presence/absence variation in *F. oxysporum* genomes without a *forma specialis* (f. sp.) annotation. (A) Gaps in annotated f. sp. *niveum* strains (**gray**) correspond to genomes originating from *Citrullus lanatus* samples (**red** arrows). (B) Gaps in annotated f. sp. *lycopersici* strains (**magenta**) are filled by genomes resulting from *Solanum lycopersicum* samples (**red** arrows) and one genome associated with a *Solanum tuberosum* sample (**red** asterisk). (C) non-annotated samples clustering together with the f. sp. *pisi* HDV247 sample.

While in the examples above all members of a *forma specialis* reside in a single cluster, other *formae speciales* are distributed in multiple clusters. In some cases, this reflects differences in disease symptoms. For example, in strawberry-infecting isolates (f. sp. *fragariae*) distinct disease phenotypes have been described. These two phenotypes, ‘yellowing’ and ‘wilting’, likely require different sets of genetic tools to successfully cause disease in the host and have traversed independent evolutionary paths to pathogenicity towards cultivated strawberry (Henry et al. 2021). In line with these previous observations, three main strawberry / berry affecting groups reside in completely different clusters. The largest group (Figure 2, bottom) is the strawberry-yellows causing clade. In contrast to the wilt-type *fragariae* isolates, the yellows isolates possess a pathogenicity chromosomal fragment “chrY”. Indeed, a yellows-specific group of putative effectors is clearly visible in these isolates. Previously, strawberry-wilting isolates have been classified into three groups based on a phylogeny of conserved core genes, where W1 and W2 consist of Australian isolates that cluster phylogenetically with strawberries-yellow isolates, but here form a separate cluster with f. sp. *mori*, affecting blackberry, and W3 consists of Spanish isolates, that are in a distinct cluster with isolates that are assigned to f. sp. *vasinfectum*. Similarly, *F. oxysporum* f. sp. *radicis-cucumerinum*, which affects multiple cucurbit crops and causes root and shoot rot rather than the wilt inducing f. *sp. cucumerinum* isolates, forms a separate, distinct cluster from other cucurbit-infecting isolates, together with two *luffae* isolates and one *lagenariae* isolate. Finally, in line with the observations by Batson et al. (2021), the two pathogenicity types that can be distinguished in f. sp. *spinaciae* also form two groups, albeit within one cluster.

Several *formae speciales* are dispersed in distinct clusters that do not correspond to known different pathogenicity types or symptoms. For example, four clusters of cucurbit-infecting strains were observed. The first is comprised of a large number of *melonis* isolates. The second contains the ‘C1’ and ‘C2’ subclades of f. sp. *melonis* identified previously (Sabahi et al., 2021), as well as isolates of the ff. spp. *momordicae, lagenariae* and *cucumerinum*. The third clade groups together representatives of f. sp. *niveum* and *cucumerinum* isolates Foc 018, 021 and 030 (van Dam et al., 2016). Like *melonis*, f. sp. *cucumerinum* is also represented in both of the main cucurbit clades in the plot. This was also observed in van Dam et al. (2016), where the abovementioned three *cucumerinum* isolates grouped away from the rest of the *cucumerinum* isolates. Similarly, f. sp. *vasinfectum* has four individual effector pattern groups. Finally, *formae speciales* that infect closely related hosts do not necessarily contain similar sets of effectors. For example, the placement of f. sp. *nicotianae* far away from other *Solanaceae* infecting isolates (*F. oxysporum* f. sp. *lycopersici, physali*) indicates different genes underlying pathogenicity of *F. oxysporum* towards these related plant species. For reference, the cluster plot including sample names, accession numbers and *formae speciales* is included in Supplementary Figure S3.

### 4.4 Core phylogeny highlights the polyphyletic nature of many *formae speciales*

To compare the clustering of effectors with the phylogenetic association of the analyzed isolates, a phylogenetic tree was generated based on core genes (Supplementary Figure S4). This tree was built based on a concatenated multiple sequence alignment of 3,037 conserved BUSCO genes, with a total length of 1,444,221 amino acid positions. Three major taxonomic clades are recognized within the *Fusarium oxysporum* species complex: clades 1, 2 and 3 (O’Donnell et al., 1998, Laurence et al., 2014). The majority (381) of the 537 *F. oxysporum* isolates fall inside main clade 2; 127 isolates belong to clade 3 and only 27 to clade 1. Two isolates do not clearly fall within one of the three clades in this tree; Fo65 (GCA_014324465.1) falls outside clade 1 and Fo24 (GCA_014337855.1) is positioned between clades 1 and 2/3, in line with the findings by Constantin et al. (2021). Many *formae speciales* clustering together in Figure 2 are polyphyletic, i.e., they occur in distinct clades within the core phylogeny. Two very pronounced examples of this include *F. oxysporum* f. sp. *lycopersici* and *F. oxysporum* f. sp. *niveum* (Supplementary Figure S4, red and blue text highlights). These two *formae speciales* are each grouped in a single cluster in Figure 2, while in the core phylogeny they are located in four and five distinct clades, respectively. Various other examples of this can be found and are supportive of HCT as the mechanism of host-specific effector gene transfer.

Interestingly, the core phylogeny (Supplementary Figure S4) shows that three of the *nicotianae* samples are phylogenetically close to a group of *lycopersici* and *physali* isolates, while their effector profiles are very different.

### 4.5 Most *SIX* genes are detected by FoEC2

In order to evaluate whether verified effectors are identified, *SIX1-SIX14* were searched for in the putative effector families produced by FoEC and FoEC2. Similar to the original pipeline, not all *SIX* genes were detected by FoEC2. *SIX5* and *SIX12* were not detected by either version of the pipeline, whereas *SIX10* was found by FoEC2 using 538 genomes, but not by FoEC or FoEC2 in the set of 59 genomes. The remaining *SIX* genes were all found at least once among the final putative effector clusters.

## 5 Discussion

### 5.1 Clustering based on putative effector profiles gives an indication of possible host species and supports horizontal chromosome transfer

In this paper, we generated the largest set of putative effectors in *F. oxysporum* to date: 1,251 effector families in 537 *F. oxysporum* genomes. It is likely that a substantial part of this number represents false positives. The pipeline allows stricter filtering by altering certain thresholds (e.g., cysteine content, protein length, etc.) while identifying putative effectors, but in this paper, settings that resulted in as few false negatives as possible were used, assessed through the detection of *SIX* genes.

The *formae speciales* clusters (Figure 2) largely align with earlier observations in other studies. The fact that various *formae speciales* are broken up into multiple clades supports the theory that pathogenicity towards a plant host sometimes evolved through multiple independent events. This seems to be the case for *F. oxysporum* f. sp. *fragariae* (three clades, Henry et al., 2021), *melonis* (three clades, Sabahi et al., 2021), *cucumerinum* (three clades) *spinaciae* (two clades, Batson et al., 2021). Isolates with a similar effector repertoire and position in the hierarchical clustering (Figure 2), but with distinct core phylogenies (Supplementary Figure S4) support the prevalence of HCT within the FOSC. HCT of (partial) pathogenicity chromosomes and thus effector genes has been experimentally shown for multiple *formae speciales*, including *lycopersici* and *radicis-cucumerinum* (van Dam et al., 2017a, Li et al., 2020, Vlaardingerbroek et al., 2016, Ma et al., 2010).

Traces of core phylogeny similarities (and thus influence of non-lineage specific candidate effector genes) can be seen in the effector clustering plot (Figure 2), but these seem to have a limited effect. For example, *F. oxysporum* f. sp. *cucumerinum* 011 and 013 (GCA_001703455.1 and GCA_001702495.1) are close to *cubense* TR4 isolates even though they do not share a related plant host. These are all core phylogeny clade 1 isolates. However, in between is a *F. oxysporum* f. sp. *koae* isolate, which is in core phylogeny clade 2.

The clustering of non-annotated isolates with isolates that do have a known *forma specialis* description (Figure 3) provides support for their hypothesized *forma specialis*. This should still be confirmed by performing virulence assays on the suspected plant host species, but this analysis provides an important clue on what host an uncharacterized isolate may cause disease on. Additionally, the results can serve to identify *forma specialis* or even subclade specific marker sequences (van Dam et al., 2018).

### 5.2 False negatives due to absence of a signal peptide prediction or large sequence diversity

Some *SIX* genes were not found among the final list of putative effectors returned by FoEC2. *SIX5* and *SIX12* have not been predicted by either version of the pipeline. In the case of *SIX12*, its product does not have a signal peptide (Schmidt et al., 2013) and is thus filtered out by the pipeline. The *SIX5* product does seem to have a signal peptide but this was not predicted by SignalP. Obviously, effectors without a signal peptide recognized by SignalP are not identified with this pipeline.

While running FoEC2 on 59 genomes, *SIX10* was found and deemed a putative effector after the first part of the pipeline, but the nhmmer hits never met the minimum query coverage of 80% due to the large sequence variation inside intronic regions. Sometimes, several of these hits were found on the same contig. To improve upon the detection of *SIX10* and similar effectors, the pipeline could consider multiple hits as a single entity when calculating the query coverage if they could potentially represent a single gene. Another solution could be to simply reduce the query coverage threshold, but that would result in a larger number of false positives.

### 5.3 The use of nhmmer increases sensitivity

One of the significant changes in FoEC2 in comparison to FoEC is the use of HMM profiles to represent clusters of putative effectors instead of the longest sequence in a cluster. HMM profiles are capable of capturing more information, keeping track of position-specific nucleotide conservation and gap or insertion frequency. The longest sequence in a cluster does not necessarily reflect information which is true for all sequences, and sequence variation is not captured when searching for presence/absence. For instance, an insertion in the longest sequence could have a negative influence on the BLAST search results. Using nhmmer is more time-consuming than blastn, but nhmmer’s increased sensitivity can identify sequences with lower similarity.

### 5.4 A more efficient and user-friendly effector detection pipeline

Previously, a few preparation steps were sometimes needed to run FoEC. One of these steps included changing FASTA file extensions to ‘.fa’ or ‘.fasta’ if ‘.fna’ was used. This was a common occurrence, given that files downloaded from NCBI typically have the ‘.fna’ extension. FoEC2 accepts multiple FASTA file extensions.

Another preparation step needed involved changing sequence headers in all genomes to a consistent format (e.g., ‘contig_1’) before running the pipeline. This naming scheme was used because FoEC stored information in the headers. FoEC2 instead uses GFF3 files and tables to store information. This eliminates the need to change the headers before a run and provides the advantage of having easily accessible information. Researchers can also visualize elements such as mimps, TIRs and putative effectors in a genome browser of their choice thanks to the new GFF3 output.

### 5.5 Conclusion

The updated FoEC2 pipeline provides an easy way for researchers to determine an initial *forma specialis* classification for *F. oxysporum* genomes and search for putative effectors within them. This can directly influence the time and resources needed to characterize newly assembled *F. oxysporum* genomes by reducing the number of experiments required to confirm host specificity. The increased run time caused by using nhmmer searching is compensated by the use of multithreading facilitated by Snakemake, the chosen framework for FoEC2. The new RShiny app provides the visualization and allows users to customize plots to suit their needs. The added MSA visualization also provides an opportunity to dive into the determined putative effector clusters. Overall, the updated FoEC2 pipeline has multiple new features contained within a modularized pipeline, better accommodating the needs of users as well as making it possible to easily expand its range of functions in the future.

## 6 Figure captions

**Supplementary Figure S1**. Examples of the FoEC2 interface. (A) ‘Data’ tab. The left sidebar ‘Input Files’ shows the three files uploaded to the app. In the main panel, the genome metadata tab is selected. Here, the metadata can be modified and saved as a CSV file. If any changes are made, the ‘Update Plot’ button can be pressed to display those changes in the ‘Plots’ tab. (B) The heatmap shows how genomes (rows) and putative effectors (columns) cluster. This interactive visualization allows users to change several parameters found in the left sidebar ‘Options’. (C) The dendrograms correspond with the distance and clustering methods selected to create the heatmap and can be downloaded as newick files. (D) The MSAs use the putative effector MSA output generated by the pipeline. Each one represents an alignment of a cluster of putative effectors detected across multiple genomes. A drop-down menu can be used to select which putative effector cluster MSA should be visualized using the BioJS MSA viewer.

**Supplementary Figure S2**. Presence (dark blue) / absence (light blue) variation of 299 putative effectors (columns) across 59 F. oxysporum genomes (rows). Genomes are annotated by *forma specialis* when possible. Putative effectors are annotated by SIX gene. Presence is determined by nhmmer, given the thresholds E-value <= 10E-10, query coverage <= 80%. Clusters were achieved with a binary distance matrix and average clustering.

**Supplementary Figure S3**. Clustering of all 538 *Fusarium* genomes including labels and at high resolution.

**Supplementary Figure S4**. Phylogenetic tree of 537 *F. oxysporum* genomes and one *F. verticillioides* genome (root) based on the concatenated alignment of 1,444,221 amino acid positions. These positions were taken from 3,037 single-copy BUSCO genes (using BUSCO lineage ‘hypocreales_odb10’) which were found in at least 98% of the genomes. *F. oxysporum* clades 1, 2 and 3 are shown in green, red and orange backgrounds, respectively. Isolates of *F. oxysporum* f. sp. *lycopersici* are highlighted with red text and *F. oxysporum* f. sp. *niveum* isolates are highlighted with blue text to illustrate their polyphyletic nature, as opposed to the single cluster for each *forma specialis* observed in Figure 2.

## Author Contributions

All authors contributed to conceiving and designing the analysis. MBG collected the data and performed the analysis. Testing of the code was performed by PvD and LF. MBG, LB and PvD wrote the manuscript.

## Acknowledgments

--

