## Supplementary figures and images for "*Fusarium oxysporum* Effector Clustering version 2 (FoEC2): an updated pipeline to infer host range"

### Figure-S1

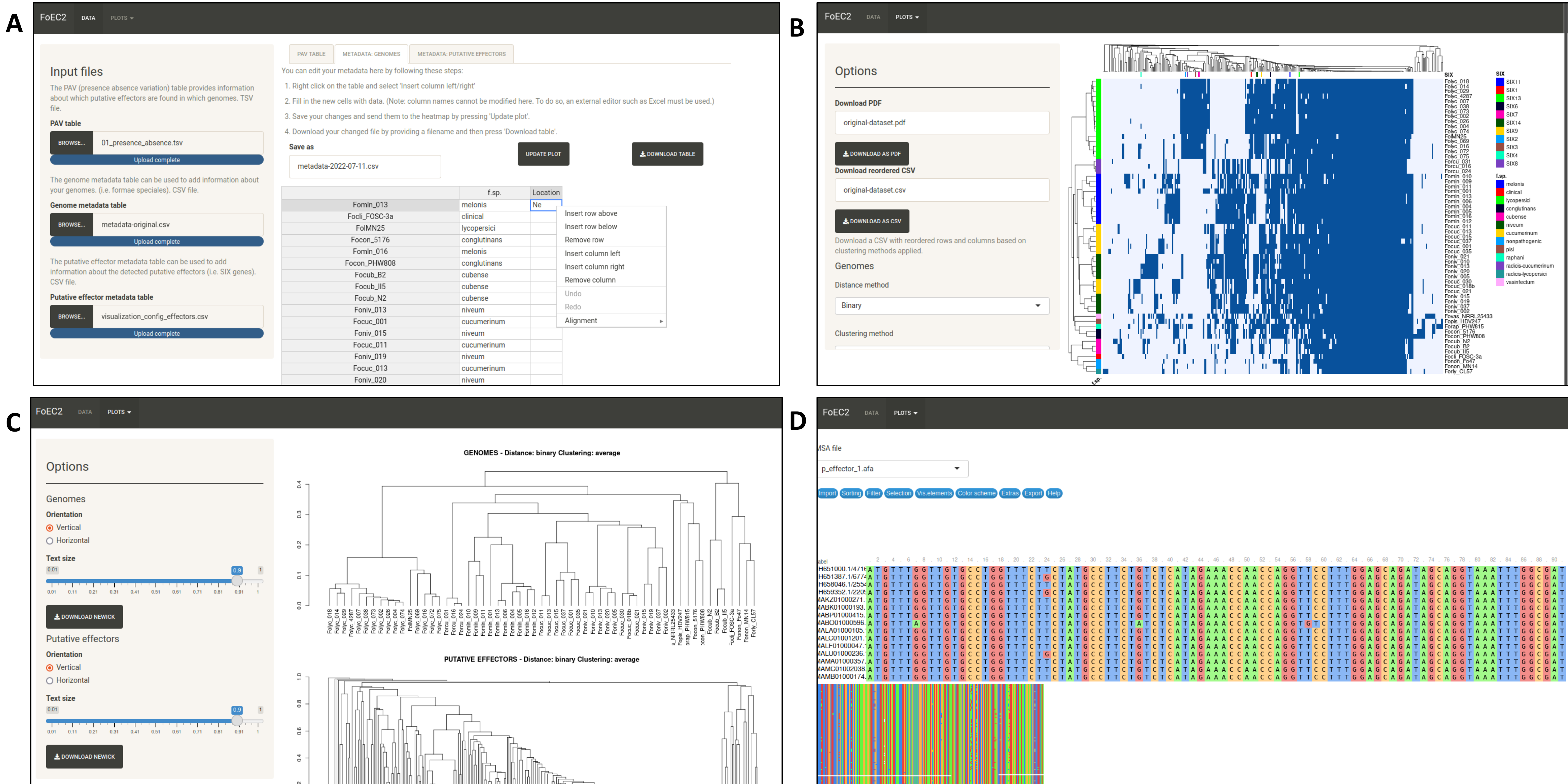

### Figure-S2

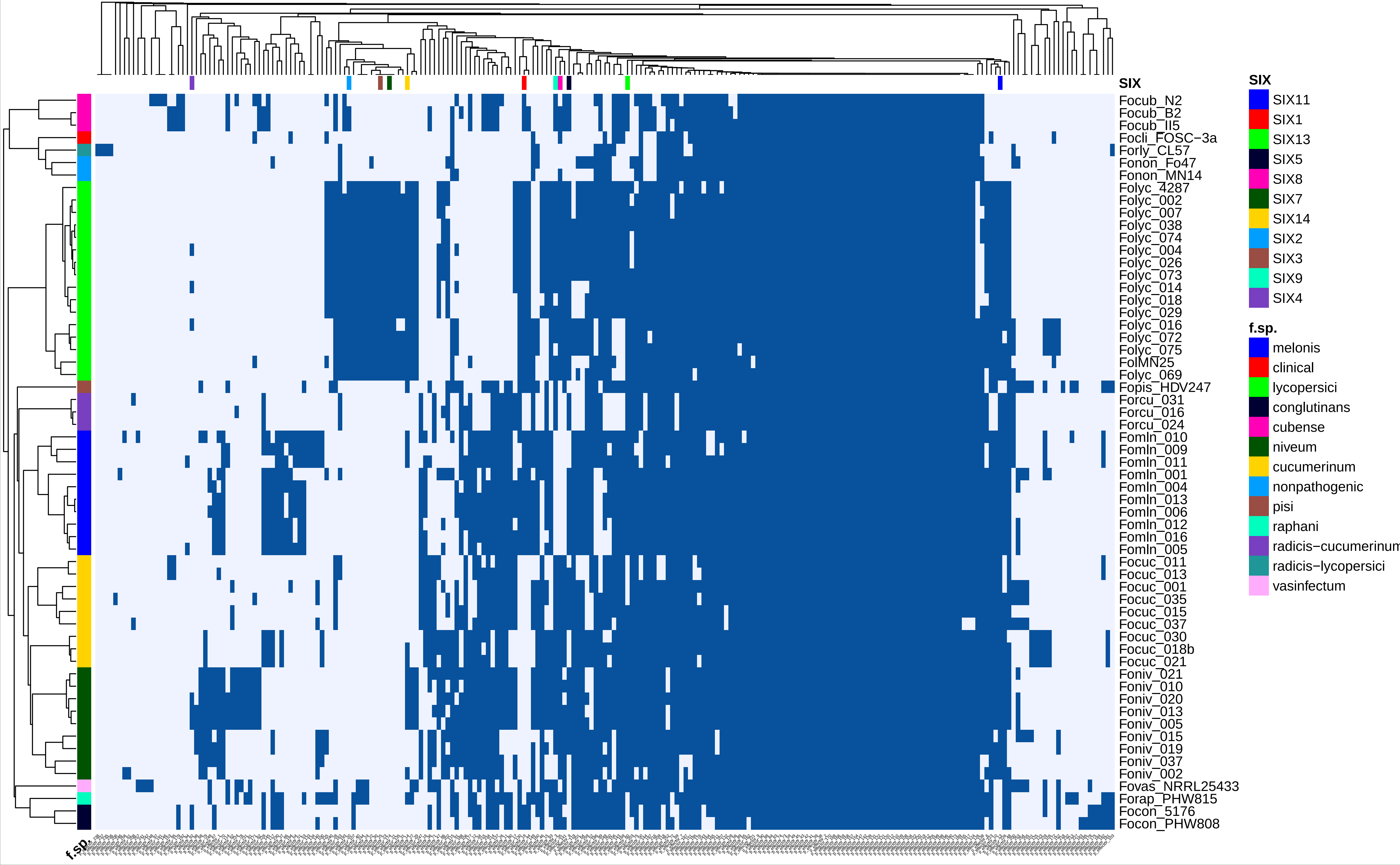

### Figure-S3

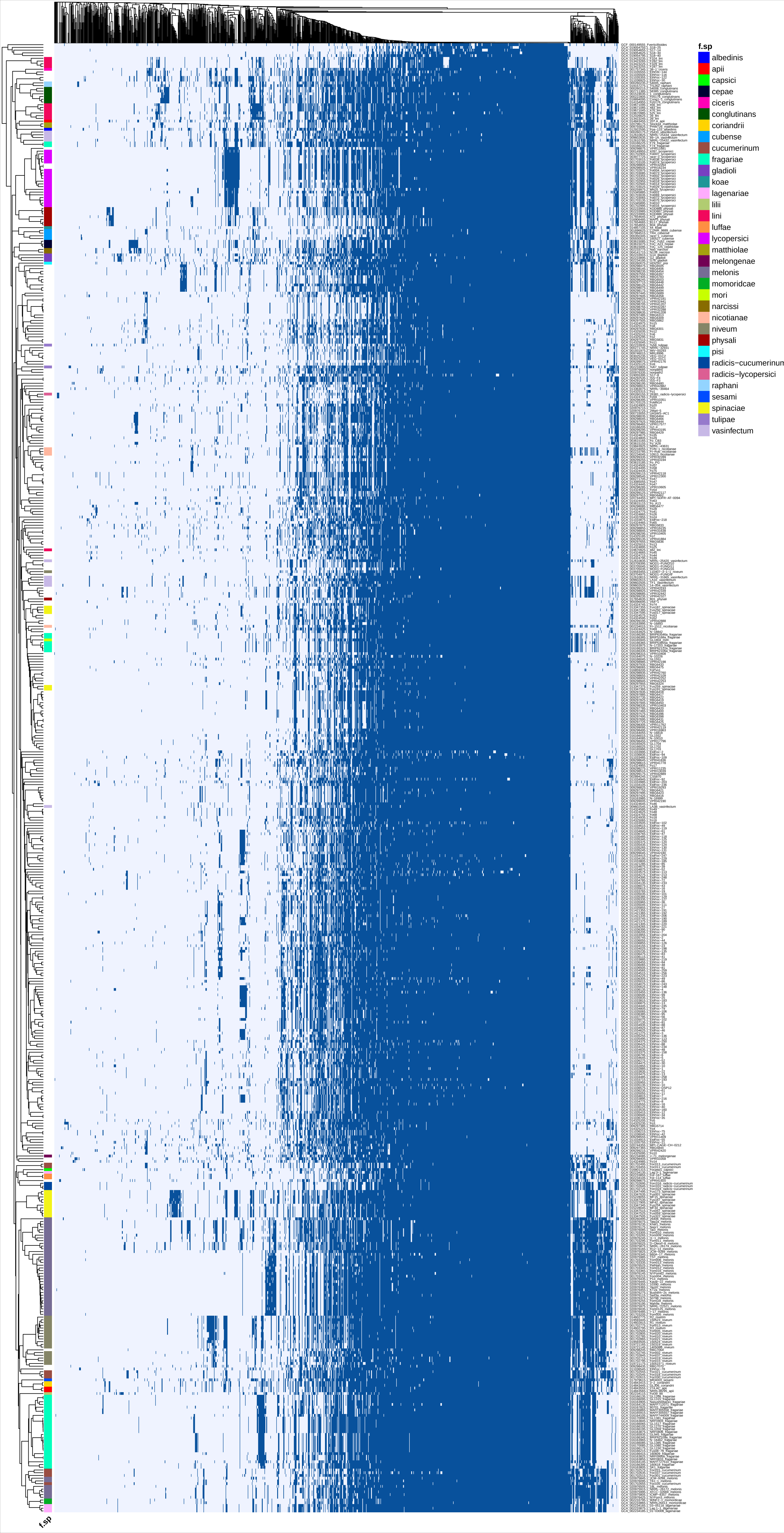

### Figure-S4

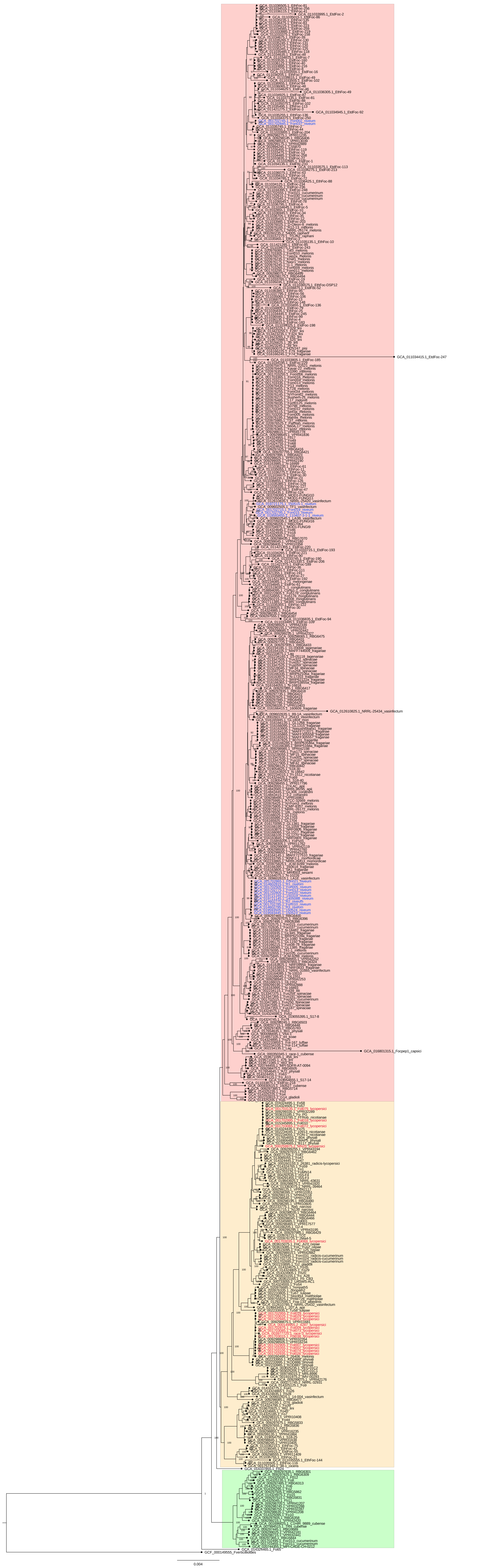
